# IGLV3-21^R110^-directed bispecific antibodies activate T cells and promote killing in a high-risk subset of chronic lymphocytic leukemia

**DOI:** 10.1101/2025.02.04.636461

**Authors:** Claudia Fischer, Simon Stücheli, Shih-Shih Chen, Johanna Nimmerfroh, Christoph Schultheiß, Corinne Widmer, Dominik Heim, Benjamin Kasenda, Jakob Passweg, Sebastian Kobold, Lukas Egli, Nicolò Coianiz, Obinna Chijioke, Nicholas Chiorazzi, Marie Follo, Heinz Läubli, Matthias Peipp, Mascha Binder

**Author notes:** **Correspondence to:** Mascha Binder, MD, Division of Medical Oncology, University Hospital Basel, Petersgraben 4, 4031 Basel, Switzerland, Phone (+) 41 79 416 5839.

## Abstract

We previously used a disease-specific B cell receptor (BCR) point mutation (IGLV3-21^R110^) for selective targeting of a poor-risk subset of chronic lymphocytic leukemia (CLL) with chimeric antigen receptor (CAR) T cells. Since CLL is a disease of the elderly and a significant fraction of patients is not able to physically tolerate CAR T cell treatment, we explored bispecific antibodies as an alternative for precision targeting of this tumor mutation. Heterodimeric IgG1-based antibodies consisting of a fragment crystallizable region (Fc) attached to either an anti-IGLV3-21^R110^ Fab or an anti-CD3 (UCHT1) single chain variable fragment (R110-bsAb) selectively killed cell lines engineered to express high levels of the neoepitope as well as primary CLL cells using healthy donor and CLL patient-derived T cells as effectors. R110-bsAb spared polyclonal human B cells (as opposed to CD19-targeting Blinatumomab) as well as CD34^+^ human stem cells. Yet, R110-bsAb induced lower T cell activation than Blinatumomab with primary CLL cells likely due to lower expression of target antigen. In vivo, R110-bsAb specifically killed IGLV3-21^R110^-expressing cell lines and CLL cells while sparing peripheral blood mononuclear cells. These findings highlight bispecific antibodies as a promising, off-the-shelf immunotherapy for high-risk CLL patients, offering selective targeting while preserving healthy B cells.

## INTRODUCTION

The treatment landscape for chronic lymphocytic leukemia (CLL) has undergone profound changes, evolving from predominantly chemotherapy and antibody combinations to the use of small molecule inhibitors targeting the B cell receptor (BCR) and BCL2 pathways (1–3). With these novel therapies, life expectancy is now approaching that of the general population (3). However, certain patient subsets do not yet experience the same benefits, highlighting the need for novel treatment options (3). These include patients from stereotypic BCR subsets, which may have a poor prognosis and derive only limited long-term benefits from current approaches, including BCR pathway-targeted therapies (4–6).

Stereotypic subsets are characterized by distinct complementarity-determining region 3 (CDR3) sequence motifs, oligomeric membrane organization, and autonomous signaling through BCR-BCR interactions (1, 7–11). Stereotypy may also extend to the BCR’s light chain (10, 12, 13). The IGLV3-21^R110^ subset is one such light chain-defined subset that typically exhibits an aggressive clinical course (11, 13–16). IGLV3-21^R110^ is expressed in 10-15% of unselected CLL patients but is overrepresented in cases requiring treatment (13, 16). Functionally, the G-to-R exchange at position 110 of the IGLV3-21 light chain, along with several conserved amino acids in the heavy chain, confers autonomous signaling capacity to the BCR by mediating self-interactions (13, 14, 16).

Since the IGLV3-21^R110^ BCR is specific to CLL and acts as a critical tumor driver, we hypothesized that targeting this receptor would spare normal B cells with a low risk of epitope escape due to its functional relevance. Additionally, the absence of persistent B cell aplasia could reduce infection-related complications and preserve vaccination responses (17). To this end, we previously developed IGLV3-21^R110^-targeted CAR T cells and demonstrated their function in vitro and in vivo, providing proof-of-concept that this targeting approach may be feasible (18).

In this study, we investigated whether our precision therapy approach could be adapted into an off-the-shelf bispecific antibody format to extend its applicability to a broader range of CLL patients, including those too frail for CAR T cell therapy.

## METHODS

### Cell lines, primary CLL and healthy donor blood cells

Cell lines (DMSZ) and IGLV3-21^R110^ or IGLV3-21^G110^ light chain expressing variants thereof were generated as previously described (18). Suspension-adapted Chinese hamster ovary (CHO)-S cells purchased from Thermo Fisher Scientific were cultivated in CD CHO-Medium (Thermo Fisher Scientific) with 1% HT Supplement (Thermo Fisher Scientific) and 1% GlutaMax-I (200 mM L-Ala-L-Gln, Thermo Fisher Scientific) added to the medium. For antibody production, they were kept in CD OptiCHO supplemented with 1% GlutaMax-I, 1% Poloxamer 188, and 1% HAT-Supplement (CHO production medium, Thermo Fisher Scientific).

Blood samples from CLL patients were collected after informed consent as approved by the ethics committees of the Universities of Hamburg–Eppendorf, Halle-Wittenberg and Basel. Peripheral blood mononuclear cells (PBMCs) were isolated by Ficoll gradient centrifugation (Cytiva). If necessary, Pan T cells, Pan B cells or CD34^+^ hematopoietic stem cells were additionally isolated via magnetic-activated cell sorting (MACS, Miltenyi Biotec). PBMCs and T cells were resuspended in FCS + 10% Dimethyl sulfoxide (DMSO) and cryopreserved in liquid nitrogen. IGLV3-21^R110^ expression was characterized by next-generation sequencing (NGS) of the light chain loci as previously described (19–28).

### Bispecific antibody constructs

The bispecific antibody construct is derived from the humanized antigen-binding fragment (Fab) of the IGLV3-21^R110^-specific antibody from AVA Lifescience GmbH (Denzlingen, Germany; patent EP 4 227 322 A1). The R110 bispecific antibody (R110-bsAb) was designed as a heterodimeric IgG1-based antibody consisting of a fragment crystallizable region (Fc) attached to either an anti-IGLV3-21^R110^ Fab or an anti-CD3 (UCHT1) single chain variable fragment (anti-CD3 scFv). Its light and heavy chain fragments were cloned into the pcDNA3.1(+) vector containing a CMV promotor for antibody expression in CHO-S cells. Knob-into-whole mutations (S354C, T366W vs. Y349C, T366S, L368A, Y407V) in the constant fragments (Fc) of its heavy chains facilitate heterodimerization (29). L234A and L235A point mutations were induced to reduce unspecific Fc-FcR interactions (30).

### Bispecific antibody production and purification

Antibody production was performed using the MaxCyte STX Scalable Transfection Systems (31). In brief, CHO-S cells were cultivated at 37°C, 6% CO_2_, 145 rpm in serum free CD-CHO medium for at least 2 weeks before being transfected via the MaxCyte STX Scalable Transfection System (MaxCyte, (32)). Transfected cells were further incubated at 32°C, 6% CO_2_, 145 rpm for 24 h before sodium butyrate (Sigma) was added. During the following days, feed stock solution, which contained 70% CHO CD Efficient Feed A Stock Solution (Thermo Fisher Scientific), 14% Yeastolate TC UF (Becton Dickinson), 3.5% GlutaMax-I(200 mM) and 12.5% Glucose (450 g/L, Sigma), was added to the cells until the cell viability dropped below 50%. Supernatant containing the antibody was filtered. For purification, antibody was isolated with the Capture Select™ CH1 Affinity Matrix (Thermo Fisher Scientific) following the manufacturer’s instructions. Multimers were excluded via size exclusion chromatography using the Äkta Chromatography System (Cytiva). The commercial bispecific antibody Blinatumomab (anti-CD19/anti-CD3, Blincyto^®^, Amgen) was used as positive control (33).

### In vitro cytotoxicity assay and cytokine quantification

For in vitro cytotoxicity assays using NALM-6 Luc (-R110), NALM-6, RAJI or OCI-LY1 (G110/R110) cells, 2×10^4^ target cells were seeded with effector cells in a 96-well plate and co-incubated in complete media (RPMI 1640 Medium, GlutaMAX™ Supplement, 100 U/mL penicillin and 100 mg/mL streptomycin, 50 µM beta-Mercaptoethanol, Thermo Fisher Scientific; 10% Human Serum) at 37°C and 5% CO_2_. For primary CLL target cells, 4×10^4^ cells were seeded. The different E:T ratios as well as the bsAb concentrations used for experiments are indicated in the respective figure descriptions. Cell lysis of target cells was determined after 24 hours of co-culture via flow cytometry using a CytoFLEX™ (Beckman Coulter). Deviating incubation periods are indicated in respective figure descriptions. CD22 PE/Dazzle 594 (HIB22), CD2 FITC (TS1/8) and CD3 APC-Cy7 (SK7, BioLegend) or CD3 FITC (SK7, BD Biosciences) as well as DAPI (Miltenyi) were used for differentiation of effector and target cells or live and dead cells. bsAb-dependent T cell activation was determined using CD8 FITC (SK1), CD25 PE (M-A251, BD Biosciences), CD8 BV650 (SK1), CD4 BV605 (SK3), CD25 BV510 (M-A251) and CD69 APC (FN50, BioLegend) antibodies. The FlowJo™ Software (v.10.10.0) was used for flow cytometric analysis. The percentage of specific killing was determined by the following formula:

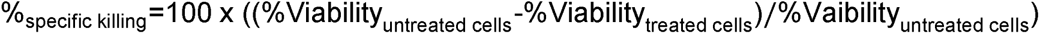

Specific target cell lysis was calculated by division of the untreated target cell lysis mean and the sample cell lysis. Fold-change of activation marker-expressing effector cells was equally calculated by the division of the percentage of untreated CD69^+^/CD25^+^ effector cells and the percentage of activation marker positive, treated samples. The amount of released IFN-γ in the supernatant of the 24 h co-culture was quantified using the LEGENDplex bead-based immunoassay (Biolegend) according to the manufacturer’s instructions. For fluorescence imaging, RAJI R110 cells were seeded at twice the usual quantity and incubated for 20 hours at 37°C and 5% CO_2_. Following Hoechst and Nile Red staining (1:100, Sigma-Aldrich), images were captured using the Zeiss LSM 710 confocal microscope and the 20x/0.8 Plan-Apochromat objective.

### In-vivo killing assays

For cell line-based xenograft assays, healthy donor T cells were activated with 25 µL/mL Immunocult Human CD3/CD28/CD2 T cell activator (STEMCELL) and expanded for 9 days in culture in Prime-XV T cell CDM supplemented with 10 ng/ml IL-7 and IL-15. A total of 10 NSG (NOD/SCID/IL2rγnull) mice were injected subcutaneously (s.c.) into the right flank with 2×10^6^ NALM-6 R110 lymphoma cells suspended in Corning® Matrigel® Matrix High Concentration Phenol-Red-Free diluted 1:1 in phenol red-free DMEM without additives. On day 7, 3×10^6^ expanded healthy donor T cells were injected intravenously either alone or with R110-bsAb (0.5 mg/kg/dose). Afterwards cells were treated biweekly with R110-bsAb for a total of 5 times. Tumor volume was measured every 2 to 3 days starting on day 10 and calculated according to the formula: D/2 x d x d, with D and d being the longest and shortest tumor diameter in mm, respectively.

Patient-derived xenograft assays were conducted as described previously (34). For each mouse, 20×10^6 PBMC were combined with 0.5×10^6^ of pre-activated T cells from the same CLL patient in a 40:1 ratio. T cell activation was achieved via CD3/CD28 dynabeads and recombinant IL-2 (R&D Systems) treatment over a period of 7 days. The cell mixture was then injected intravenously in a total of 15 NSG mice. After 10 days, the mice were divided equally into three groups (n = 5 per group) and treated intravenously with either PBS, 0.25 µg/g R110-bsAb or 0.25 µg/g Blinatumomab. The bispecific antibodies were further readministered biweekly. After three weeks, the mice were sacrificed and the spleens were harvested. T and B cell populations were quantified by flow cytometry using anti-human CD19 PE-CF594 (HIB19) and anti-human CD3 PE-Cy7 antibodies (SK7, BD Biosciences). Animal studies were performed in compliance with experimental protocols approved by the Institutional Animal Care and Use Committee (IACUC) of the Feinstein Institute for Medical Research.

Lastly, the effect of R110-bsAb on healthy, polyclonal PBMC, was analyzed using 3–5 month old NFA2 (NOD.Cg-*Rag1*^tm1Mom^ *Flt3*^tm1Irl^ *Mcph1*^Tg(HLA-A2.1)1Enge^ *Il2rg*^tm1Wjl^/J) mice that were injected intraperitoneally (i.p.) with 2.5□×□10^6^ PKH26-labelled PBMCs originating from two different donors, either alone or in the presence of 0.25µg/g of R110-bsAb or Blinatumomab (in total n□=□18). NFA2 mice were housed under a 12□h light/12□h dark cycle (lights on: 6□am, lights off: 6□pm) at temperatures from 21–24□°C with 35–70% humidity. After 16 hours, the mice were sacrificed, and peritoneal cells were collected via lavage and analyzed by flow cytometry. Mice in which PBMC cells could not be detected by flow cytometry were excluded from the dataset to control for unsuccessful i.p. injections. The antibodies anti-human CD19-BUV661 (HIB19, BD Biosciences) and Zombie NIR™□ Fixable Viability Kit (Biolegend) were used for the analysis.

### Statistical analysis

Ordinary one-way Analysis of Variance (ANOVA) or 2way ANOVA combined with a Šidák’s multiple comparisons testing was used for the comparison of cell lysis target cells and activation marker expression on effector cells in the absence or presence of bsAb. Non-linear regression models were further utilized to highlight the dose-dependent decrease of target cell viability and increase of T cell activation marker expressing effector cells. For statistical analyses, the GraphPad Prism software (v10.3.0 (461)) was used.

## RESULTS

### Anti-IGLV3-21^R110^ bispecific antibodies mediate epitope-selective tumor cell lysis in vitro

For the treatment of CLL patients with the IGLV3-21^R110^ light chain mutation (R110), we developed a bispecific antibody construct containing a R110-specific binding moiety coupled to the anti-CD3 domain UCHT1 (Fig. 1A).

**Fig. 1.**
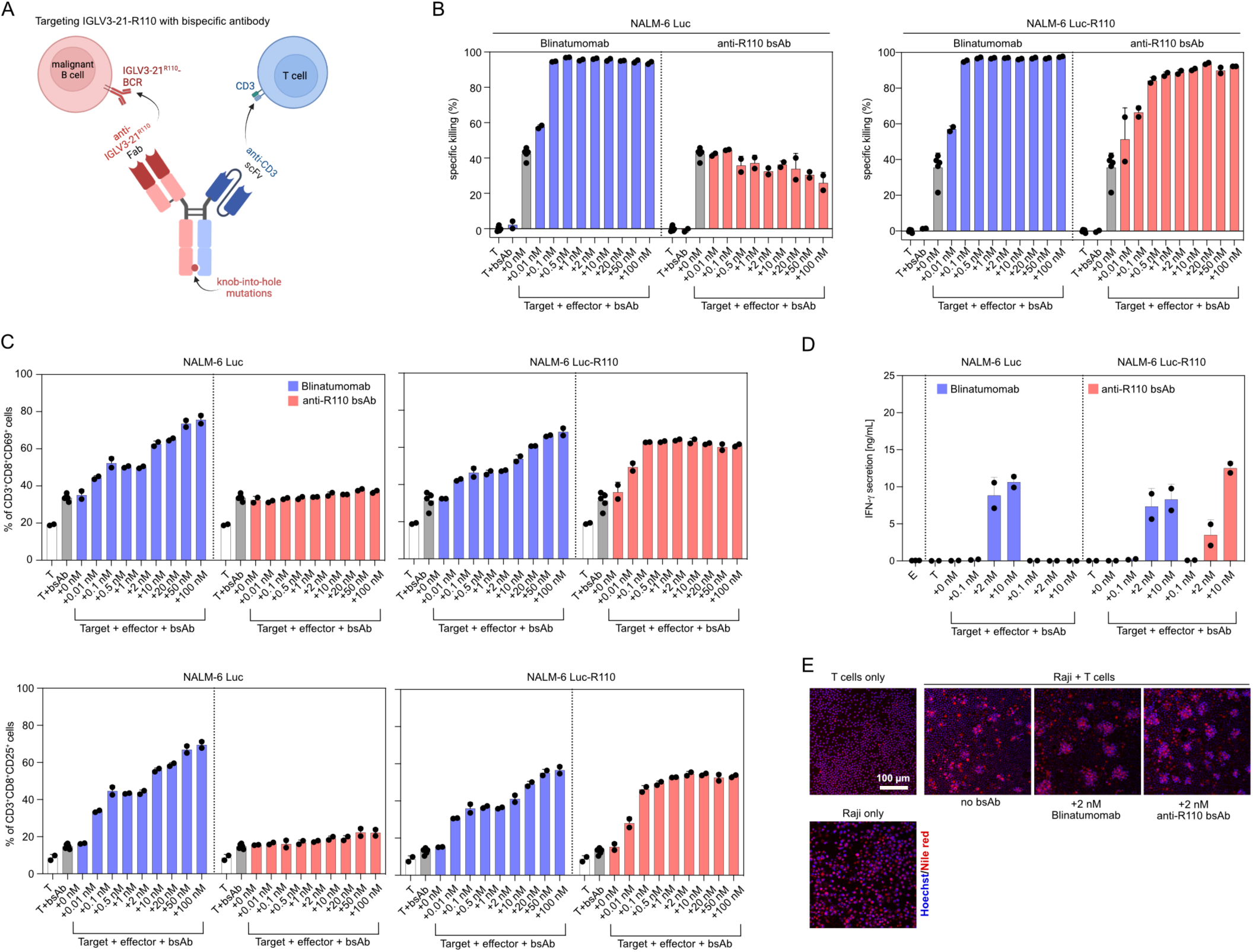
Precision targeting the IGLV3-21^R110^ neoepitope with bispecific antibodies. **A** Schematic depiction of the design and mechanism of action of R110-bsAb. Created with BioRender.com. **B** Specific killing of NALM-6 Luc and NALM-6 Luc-R110 target cells via bispecific antibody T cell engagement with an effector to target (E:T) ratio of 5:1 after 24 h. Healthy Donor (HD) T cells were used as effector cells with Blinatumomab serving as a control. Killing was normalized to the cell viability of target cells in absence of effector cells or bispecific antibodies. **C** Expression of activation markers CD69 and CD25 on CD8^+^ HD T cells after the 24 h co-culture with NALM-6 Luc or NALM-6 Luc-R110 cells (E:T = 5:1) and a non-serial dilution of bispecific antibodies Blinatumomab and R110-bsAb. Each bar plot represents the mean of at least two technical replicates with error bars as standard deviation (SD). **D** IFN-γ secretion in supernatants harvested after 24 h of co-culture with healthy donor T cells and indicated target cells. **E** Confocal microscopy pictures representative for target-effector cell engagement after 24 h incubation with 2 nM Blinatumomab or R110-bsAb stained with Hoechst for cell nucleus and Nile Red for cytoplasm. T: target, E: effector

Cell lines expressing a hybrid BCR containing the IGLV3-21^R110^ or a corresponding wild-type IGLV3-21^G110^ light chain were generated as described in a previous project (18). The conditions for co-culture experiments were set up using the NALM-6 Luc cell line as target cells with different ratios of healthy donor T cells as effector cells (effector to target ratios; E:T) and the anti-CD19 bispecific T cell engager Blinatumomab as positive control at the previously established concentration of 2 nM (Suppl. Fig. 1A)(35, 36). In addition, we used primary R110-negative CLL cells as target cells (Suppl. Fig. 1B). In both settings, an E:T ratio of 5:1 appeared to allow for efficient lysis of both the target cell line and primary CLL cells.

Co-culture of NALM-6 Luc R110 cells with healthy donor T cells and increasing concentrations of R110-bsAb showed increasing levels of epitope-selective lysis of the NALM-6 Luc R110 cell line, while control NALM-6 Luc cells were unaffected (Fig. 1B). Blinatumomab equally lysed both CD19-positive cell lines independently of the R110 epitope (Fig. 1B). Without the addition of bispecific antibodies, only baseline levels of killing were observed (Fig. 1B). If no effector cells were present, no specific killing above baseline occurred, suggesting that R110-bsAb killing was mediated by engagement of effector T cells (Fig. 1B). Cell lysis was accompanied by expression of CD25 and CD69 on the activated CD8^+^ T cells (Fig. 1C).

These results were reproducible with alternative B cell lines transduced to express the R110-epitope such as NALM-6 (without Luciferase), OCI-LY1 and RAJI (Supplementary Fig. 2A, B). To confirm specificity for the R110 point mutation, we included for each cell line a variant expressing the IGLV3-21 light chain in wild-type configuration (G110). As expected, these wild-type variants were unaffected by R110-bsAb treatment (Supplementary Fig. 2A, B). The mutation-specific pattern was also observed if IFN-γ secretion was used as a read-out (Fig. 1D). Since the RAJI cell line transduced to express the R110 neoepitope showed lowest killing rates by flow cytometry (Supplementary Fig. 2A), we wished to further corroborate our findings by confocal microscopy. Also RAJI R110 showed extensive cluster formation upon R110-bsAb treatment suggesting sufficient T cell engagement (Fig. 1E).

### T cells lyse primary CLL cells in the presence of R110-directed bispecific antibodies

We chose a CLL case with known R110-expression and a R110-negative case to test our R110-bsAb relative to CD19-directed Blinatumomab in a setting of primary human CLL cells. R110 epitope-specific patterns of cell lysis were observed with primary human CLL cells as targets using the lysis assay described above (Fig. 2A). Blinatumomab lysed primary CLL cells independently of R110 status (Fig. 2A). R110-bsAb and Blinatumomab dosing required for optimal lysis was higher in this model using primary CLL cells as compared to the cell line model. This was also reproducible when testing the R110-positive cells from a second patient (Supplementary Fig. 3). T cell activation accompanied the observed effects, but Blinatumomab more potently induced T cell activation than R110-bsAb (Fig. 2B, C). Comparing CD4 with CD8, we noted very similar activation patterns (Fig. 2B, C). Since T cell activation by the two T cell engaging antibodies was equal in the cell line model with high expression of the target antigens CD19 and R110, the differences in the assays with primary cells were interpreted to be related to lower R110 antigen density in CLL as previously shown (18).

**Fig. 2.**
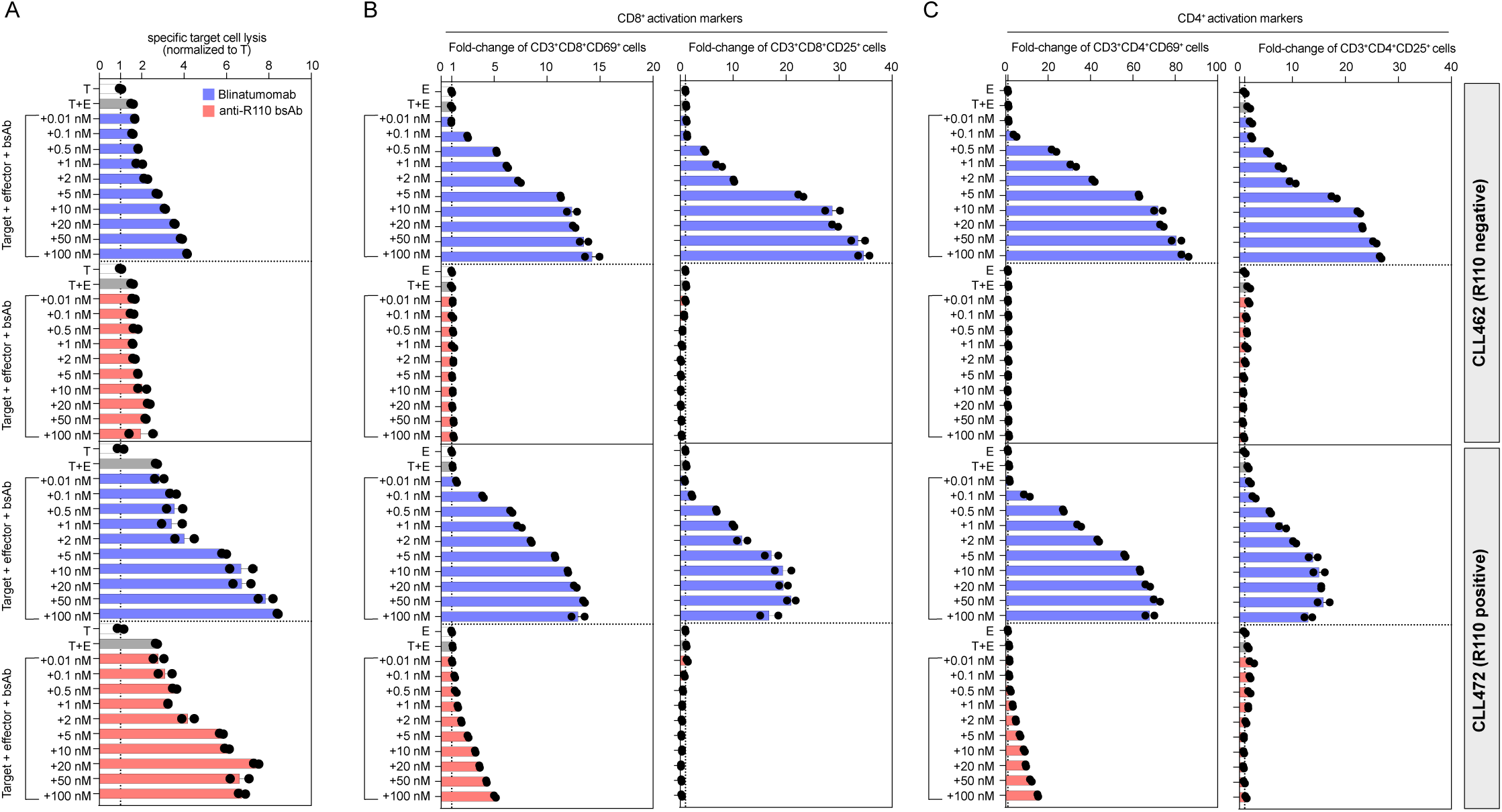
Specificity and activity of R110-bsAb against primary CLL cells. **A** Specific target cell lysis of primary target cells CLL462 (R110 negative) and CLL472 (R110 positive) after 24 h of co-culture with HD T cells. An E:T ratio of 5:1 was used in combination with a non-serial dilution of R110-bsAb or Blinatumomab. Cell lysis was normalized to the cell lysis of target cells without effector cells or bispecific antibody. **B** Fold-change of CD69 and CD25 activation marker expressing CD8^+^ HD T cells after a 24 h co-culture with primary CLL samples (E:T = 5:1) and a non-serial dilution of the bispecific antibodies. The expression was normalized to the expression of activation markers on effector cells in absence of target cells or antibodies. **C** Fold-change of CD69 and CD25 activation marker expressing CD4^+^ HD T cells after a 24 h co-culture with primary CLL samples (E:T = 5:1) and a non-serial dilution of bispecific antibodies Blinatumomab and R110-bsAb. The percentage of activated cells was normalized to the percentage of effector cells in absence of target cells or antibodies. Each bar plot represents the mean of two technical replicates with error bars as SD. T: target, E: effector.

### R110-directed bispecific antibodies spare polyclonal human B cells, peripheral blood mononuclear cells and hematopoietic stem cells

T cells in the presence of R110-bsAb did not lyse and were not activated by polyclonal human B cells at an E:T ratio of 5:1 (BC, Fig. 3A, B, C). In conditions with Blinatumomab, cell lysis and T cell activation were observed with polyclonal human B cells as target cells as expected (Fig. 3A, B, C).

**Fig. 3.**
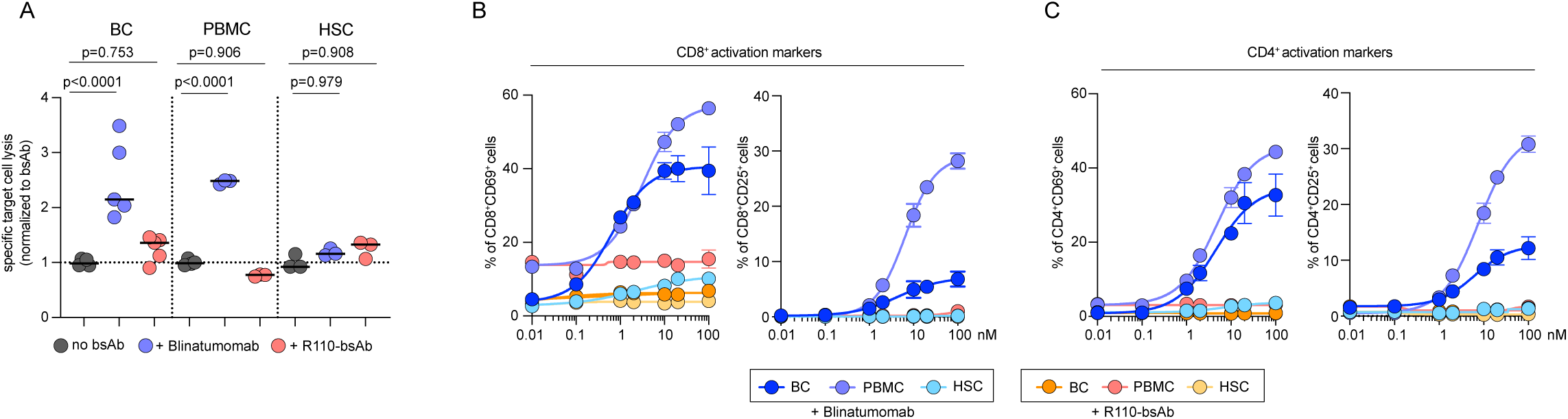
Absence of cytotoxic effects of R110-bsAb on the healthy immune cell repertoire and hematopoietic stem cells. **A** Specific cell lysis of healthy B cells (BC, PBMC) or hematopoietic stem cells (HSC) after 24 h of co-culture with HD T cells in an either allogenic (E:T = 5:1) or autologous (E:T = 9:1) system. For the allogenic systems, target cells were isolated from healthy PBMC and incubated with sorted effector cells and 100 nM Blinatumomab or R110-bsAb. In the autologous system, PBMC were incubated directly after the addition of 100 nM of bispecific antibodies. Cell Lysis was normalized to the baseline cell lysis without bispecific antibody. Each dot represents a technical replicate. **B, C** Percentage of activation marker expressing CD8^+^ (B) or CD4^+^ (C) T cells after 24 h co-culture of HD T cells cultured with BC, HSC (E:T = 5:1) or PBMC (E:T = 9:1) and a non-serial dilution of Blinatumomab or R110-bsAb. Non-linear regression analyses were performed to highlight the dose-dependent upregulation of T cell activation due to T cell engagement. Each dot represents the mean of at least three technical replicates with error bars as SD. For statistical analysis, an ordinary one-way Analysis of Variance (ANOVA) combined with a Šidák’s multiple comparisons test was used. T: target, E: effector, BC: B cells, PBMC: peripheral blood mononuclear cells, HSC: hematopoietic stem cells.

To explore B cell lysis in a more natural setting, we used healthy donor derived peripheral blood mononuclear cells (PBMC) and treated them with R110-bsAb. The natural E:T ratio in these samples was 9:1 and therefore even higher than in the previous experiments. T cells in the presence of R110-bsAb did not lyse and were not activated by PBMC, while with Blinatumomab lysis and activation were observed (Fig. 3A, B, C).

Since the R110-epitope is tumor-specific, we did not expect any reactivity with normal tissues. To explicitly rule out stem cell toxicity, we included hematopoietic CD34-positive stem cells (HSC) in our co-culture experiment. CD19-negative HSC were isolated from a leukapheresis product of a human donor by CD34 sorting. As expected, no killing or activation of T cells was observed with R110-bsAb (Fig. 3A, B, C).

### CLL patient-derived T cells lyse target cells in the presence of R110-directed bispecific antibodies

To better simulate the patient setting, we next asked if this targeting principle is also applicable to primary CLL T cells. We, therefore, isolated T cells from CLL patients and used them as effector cells in co-culture assays using the OCI-LY1 R110 model system. Indeed, the epitope-specific patterns of cell lysis were equally observed with primary CLL-derived T cells as effector cells using the lysis and activation assays described above (Fig. 4A, B, C). Blinatumomab showed epitope-independent killing with primary CLL-derived T cells in this cell line model (Fig. 4A, B, C).

**Fig. 4.**
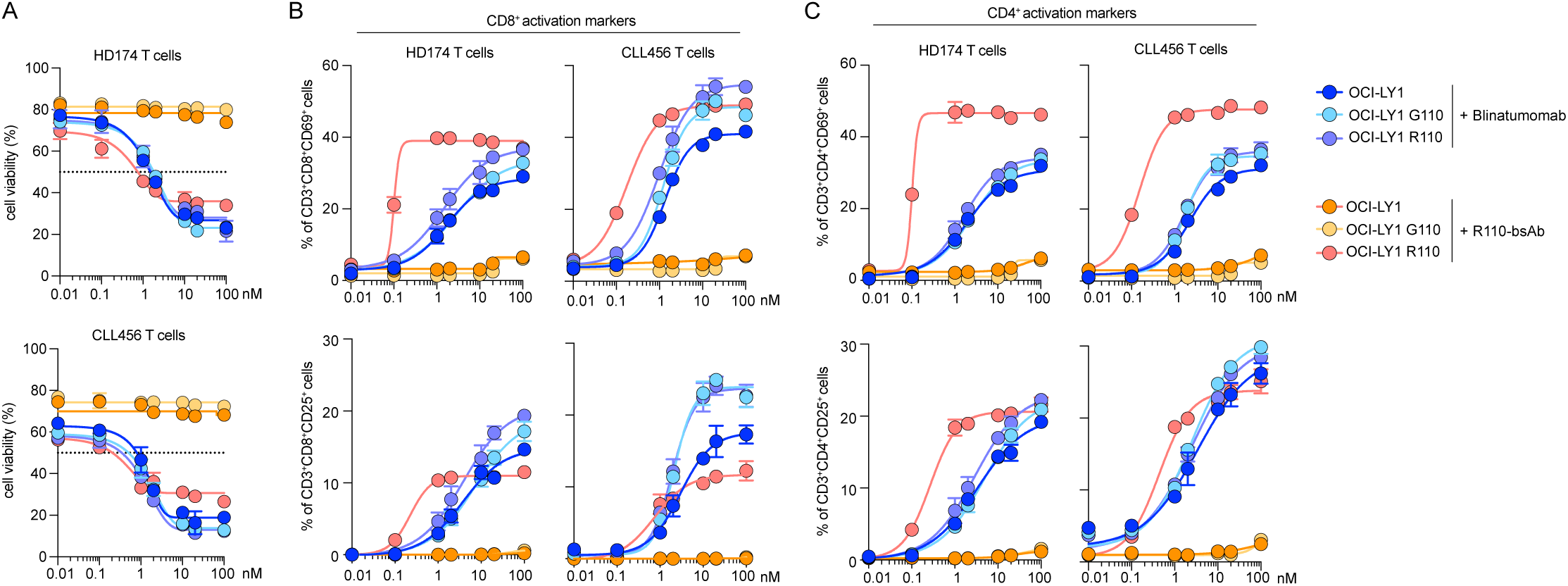
Comparison of R110-bsAb mediated cytotoxicity and activation of healthy donor or CLL patient-derived T cells. **A** Cell viability of OCI-LY1, OCI-LY1-G110 and OCI-LY1-R110 cells after 24 h of co-culture with either HD T cells or CLL T cells (E:T = 5:1) in combination with a non-serial dilution of Blinatumomab or R110-bsAb. **B** Expression of activation markers CD69 and CD25 on CD8^+^ HD or CLL T cells after 24 h co-culturing with target cells and a non-linear dilution of bispecific antibodies. **C** Expression of activation markers CD69 and CD25 on CD4^+^ HD or CLL T cells after 24 h co-culturing with target cells and a non-linear dilution of bispecific antibodies. Non-linear regression analyses were performed to demonstrate the dose-response of target cell viability and effector cell activation towards bispecific antibody treatment. Each data point represents the mean of two technical replicates (four for 2 nM) with error bars as SD.

### R110-directed bispecific antibodies are efficacious in xenograft IGLV3-21^R110^-models

Next, we evaluated the in vivo efficacy of our R110-bsAb in three different mouse models. We engrafted NSG mice with NALM-6 R110 cells and administered repeated treatments of the R110-bsAb, while untreated mice served as controls (Fig. 5A). Monitoring tumor growth over time, we observed exponential tumor growth in mice without treatment starting 20 days post tumor injection (Fig. 5B). In contrast, tumor growth was effectively suppressed in mice treated with the R110-bsAb (Fig. 5B).

**Fig. 5.**
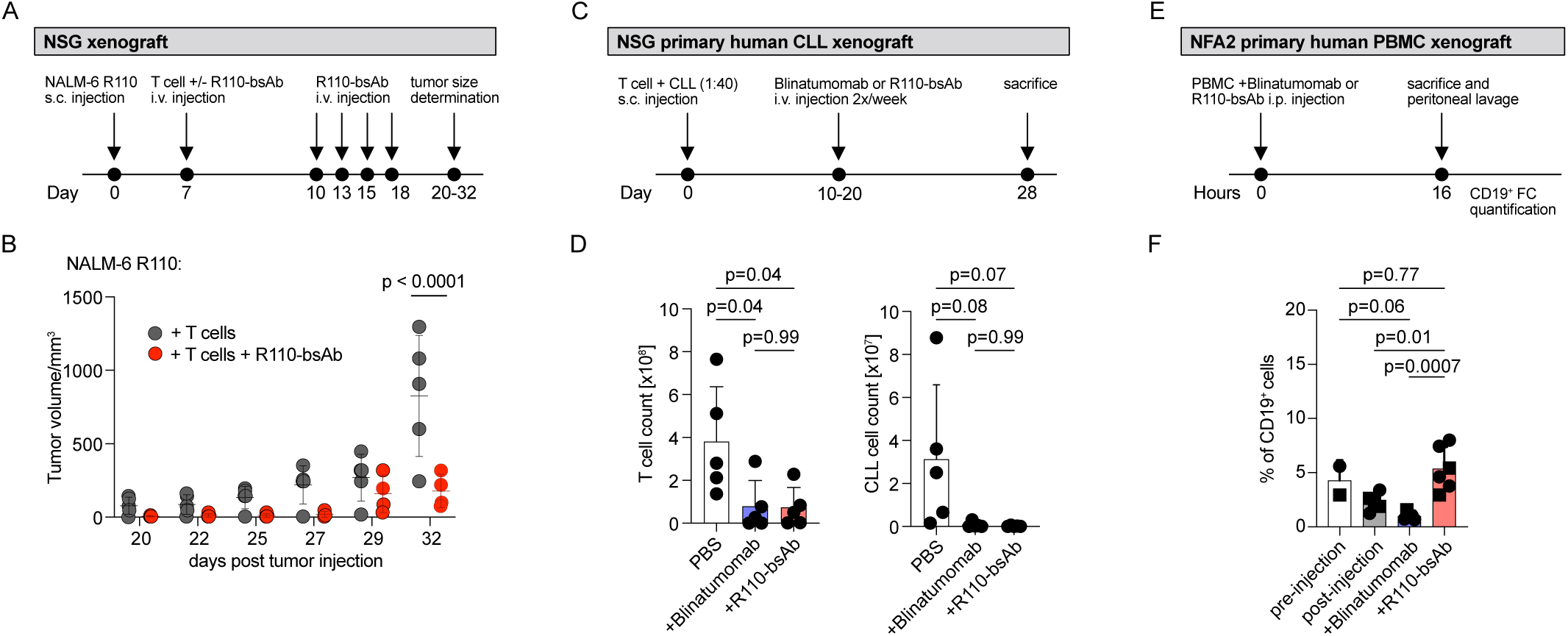
In vivo activity of R110-bsAb. **A** Workflow of the NALM-6 R110 cell line xenograft model. **B** Growth of NALM-6 R110 tumor cells subcutaneously engrafted in NSG mice untreated or treated with R110-bsAb every 2 to 3 days for 3 weeks. Engrafted mice treated only with T cells served as negative controls. Each data point represents one mouse with the mean tumor volume and error bars as SD. One outlier was identified by Grubbs’Test and removed from the analysis. Statistical analysis was performed using 2way ANOVA combined with a Šidák’s multiple comparisons test. **C** Workflow of the primary human CLL xenograft model. **D** Percentage of human CD3 and CD19 positive cells derived from spleens of NSG mice engrafted with primary CLL derived from the IGLV3-21^R110^-positive CLL donor CLL472. 20 million CLL PBMCs were i.v. injected together with 0.5 million of autologous, activated T cells. Starting on day 10, mice were treated with R110-bsAb, Blinatumomab or PBS twice a week before being sacrificed after 3 weeks. Each data point represents one mouse with error bars as SD. For statistical analysis, an ordinary one-way ANOVA with Tukey’s multiple comparisons test was used. **E** Workflow of the primary human PBMC xenograft model. **F** Percentage of human CD19 positive cells in NFA2 mice injected i.p with healthy, polyclonal PBMC +/- R110-bsAb or Blinatumomab. Mice were sacrificed after 16 hours and cells harvested from the peritoneum were analyzed via flow cytometry. Each dot presents one mouse injected with PBMC derived from one donor and each square represents one mouse injected with PBMC derived from another donor with error bars as SD. Statistical analysis was performed using the ordinary one-way ANOVA paired with Tukey’s multiple comparisons test. PBMC: peripheral blood mononuclear cells, s.c.: subcutaneous, i.v.: intravenous, i.p.: intraperitoneal.

To further validate the therapeutic efficacy of R110-bsAb, a human CLL xenograft mouse model was generated from R110-positive patient CLL472. NSG mice were injected intravenously with patient CLL472-derived B and T cells in a 40:1 ratio and treated with R110-bsAb or Blinatumomab biweekly for 3 weeks starting 10 days post CLL cell injection. (Fig. 5C). After sacrificing the mice, the distribution of human CD3^+^ and CD19^+^ cells in the mouse spleen was analyzed by flow cytometry. We observed a nearly complete clearance of B cells form the spleen of R110-bsAb and Blinatumomab treated mice (Fig. 5D). Mice treated with PBS retained high levels of B cells. Importantly, there was no significant difference between the CD3^+^ T cell counts after R110-bsAb and Blinatumomab administration (Fig. 5D).

Lastly, injecting NFA2 mice with human healthy donor polyclonal PBMC and our R110-bsAb revealed no significant reduction of the B cell population compared to the non-injected mice (Fig. 5E, F). In contrast, Blinatumomab treatment led to a significant decrease in B cell numbers compared to preinjection levels or R110-bsAb treatment (Fig. 5F).

## DISCUSSION

Cell-based and bispecific antibody-based immunotherapies have become essential treatment options for patients with B cell lymphomas, achieving long-term remission in many patients (37–44). However, in chronic lymphocytic leukemia (CLL), these therapies are not as widely used as in other lymphomas (45). Currently, only CD19 CAR T cells are approved in the U.S. for CLL, with no bispecific antibody therapies yet approved (41, 45). One challenge in using these treatments is the eradication of the entire B cell lineage, potentially leading to infectious complications and lack of response to vaccination (17, 46). Since CLL patients are often elderly and frail, the applicability of CAR T cell therapy may also be somewhat hampered due to generally more severe side effects compared to bispecific antibodies (47–49). Consequently, there is a need for more targeted and tolerable therapeutic approaches especially for patients with high-risk disease.

To address these challenges, we have developed a bispecific T cell engager that targets a recurrent oncogenic point mutation in the BCR light chain of malignant CLL cells. Previously, we demonstrated proof-of-principle with a CAR T cell therapy targeting the same epitope (18). The data presented here show that this bispecific approach is effective, even when using CLL-derived T cells as effector cells. However, T cell activation was not as robust in vitro as with a CD19-directed bispecific T cell engager, likely due to the reduced membrane expression of our target compared to CD19 (50). Importantly, however, our construct selectively spared healthy B cells, similar to the precision targeting seen in our CAR T cell approach (18).

Our research contributes to the broader effort of developing immunotherapies that target restricted, ideally tumor-specific, rather than lineage-specific, surface molecules. Several studies have explored this direction, notably the application of similar concepts to target clonotypic T cell receptors (TCR) in T cell lymphomas with antibody-drug conjugates (51) or bispecific antibodies (52). However, T cell lymphomas are less clonal, with approximately 50% of cases being oligoclonal for the TCR, which limits the applicability of this strategy (53, 54).

One of the major challenges in CLL is T cell dysfunction, which may limit the efficacy of such therapies (48, 49, 55). In our study, we observed little to no reduction in the potency of T cells recruited to lyse target cells when using patient-derived effector cells. Nevertheless, the efficacy of these T cells may vary between individual patients and at different stages of treatment. Before progressing to clinical trials, it will be important to conduct repeated testing with a variety of CLL T cell donors at different disease stages to better understand the factors influencing treatment efficacy. Given that T cell dysfunction worsens over time in these patients (55), it may be advisable to test these strategies early in the course of high-risk IGLV3-21^R110^ disease.

In summary, we provide proof-of-concept for a mutation-targeted bispecific antibody approach in CLL, which warrants further study.

## DATA AVAILABILITY

Sequences of the humanized anti-R110 Fab are publicly available (EP 4 227 322 A1).

## ACKNOWLEDGEMENTS

We acknowledge Anja Muskulus for the production of the R110-bsAb as well as the flow cytometry core facility at the Department of Biomedicine, Mihaela Barbu-Stevanovic, Cécile Cumin and Morgane Hilpert, for assistance with the generation of flow cytometry data. This work was supported by the Swiss National Science Foundation (SNSF) [10.001.762] (to M.B.).

## AUTHOR CONTRIBUTIONS

Idea & design of research project: MB, MP; Supply of critical material (e.g. patient material, mouse models, cohorts): MP, SK, CW, JP; Establishment of Methods: MP, CS, CF; Experimental work: CF, SS; Analysis and interpretation of primary data: MB, CF, CS; Drafting of manuscript: MB, CF. All authors reviewed and revised the manuscript.

