## Supplemental_Material for "IGLV3-21^R110^-directed bispecific antibodies activate T cells and promote killing in a high-risk subset of chronic lymphocytic leukemia"

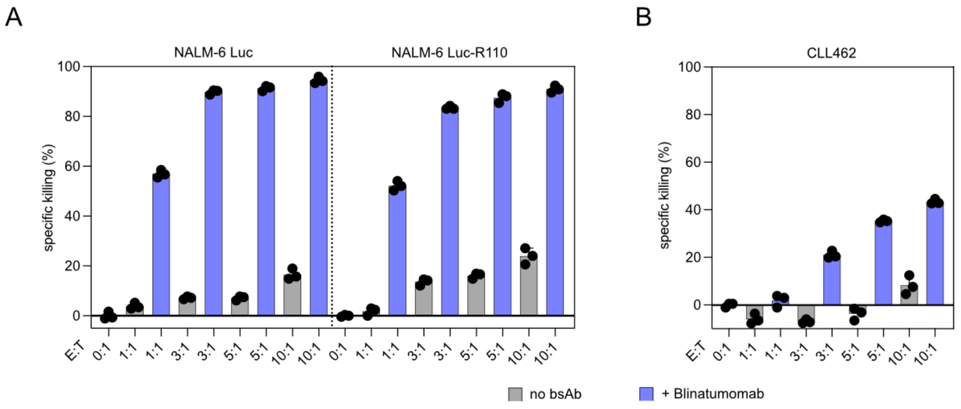


**Supplementary Figure 1. Determination of suitable E:T ratios for co-culture experiments. A** Specific killing of NALM-6 Luc and NALM-6 Luc-R110 cells incubated at different E:T ratios (0:1, 1:1, 3:1, 5:1, 10:1) with healthy donor (HD) T cells and 2 nM Blinatumomab. **B** Specific killing of primary CLL cells with HD T cells in different E:T ratios (0:1, 1:1, 3:1, 5:1, 10:1) and 5 nM of Blinatumomab. Each bar plot represents the mean of three technical replicates with error bars as SD. E: effector cells, T: target cells, bsAb: bispecific antibody.

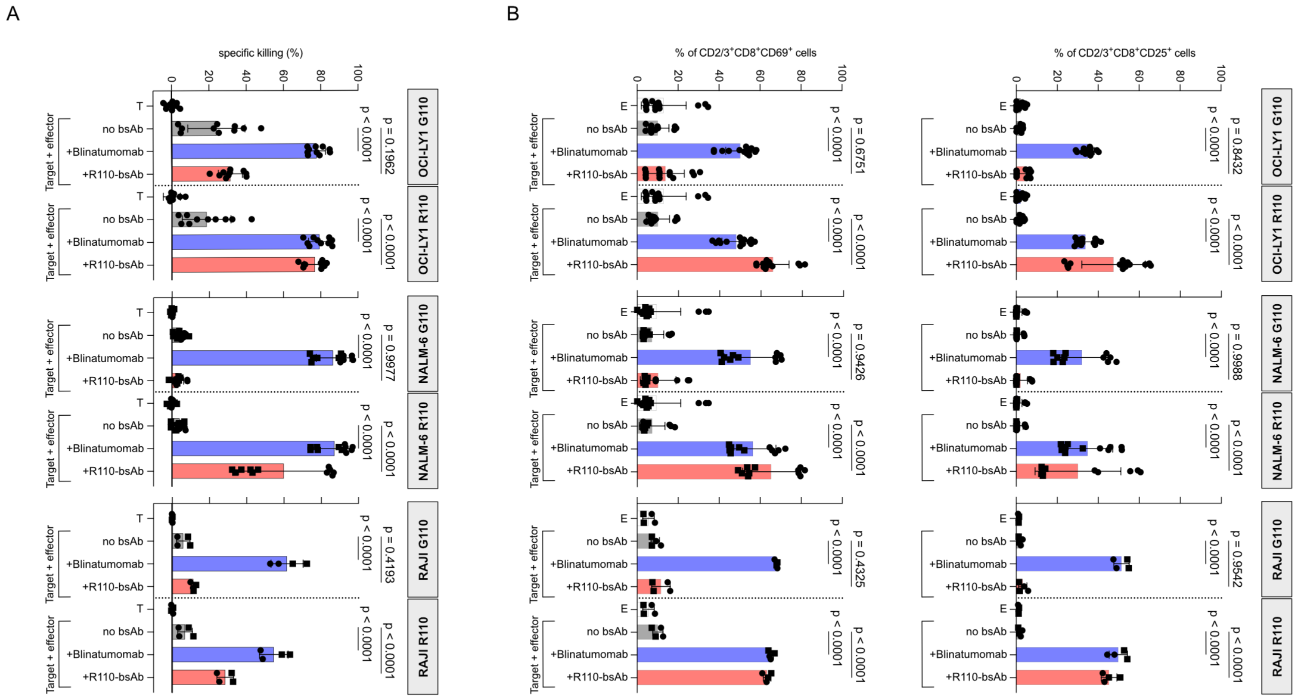


**Supplementary Figure 2. Efficacy of R110-bsAb in R110 positive cell line models. A** Cell viability of OCI-LY1 G110/R110, NALM-6 G110/R110 and RAJI G110/R110 cells incubated for 24 h with healthy donor (HD) T cells in a 5:1 ratio and 2 nM bispecific antibodies. **B** Percentage of activation marker expressing CD8⁺ HD T cells after 24 h of co-culture with different target cells treated with 2 nM Blinatumomab or R110-bsAb. Dots and squares are technical replicates representative for two different healthy donors used derived from N = 4 (OCI-LY1 G110/R110), N = 3 (NALM-6 G110/R110) or N = 1 (RAJI G110/R110) independent experiments and error bars as SD. Statistical significances were determined by ordinary one-way ANOVA combined with a Šidák`s multiple comparisons test. T: target, E: effector, bsAb: bispecific antibody.

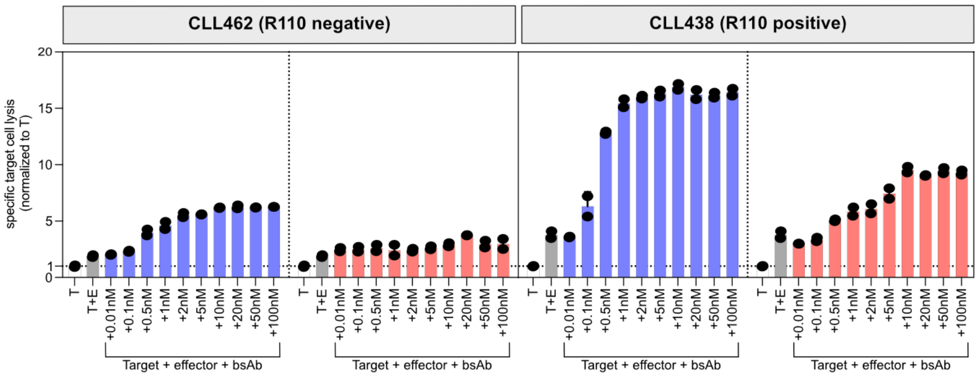


**Supplementary Figure 3. Efficacy of R110-bsAb in primary CLL.** Specific cell lysis of CLL462 and CL438 target cells with healthy donor (HD) T cells in a 5:1 E:T ratio incubated for 48 h with a non-serial dilution of Blinatumomab or R110-bsAb. Cell Lysis was normalized to the cell lysis of target cells without effector cells or bispecific antibody. Each bar plot represents the mean of two technical replicates with error bars as SD. T: target, E: effector, bsAb: bispecific antibody.
